# Massive solubility changes of neuronal proteins upon simulated traumatic brain injury reveal the role of shockwave in irreversible damage

**DOI:** 10.1101/2022.12.31.522370

**Authors:** Amir Ata Saei, Hassan Gharibi, Hezheng Lyu, Brady Nilsson, Maryam Jafari, Hans von Holst, Roman A. Zubarev

## Abstract

The immediate molecular consequences of traumatic brain injuries or TBI are poorly understood. Here, we simulated TBI using an innovative laboratory apparatus that employs a 5.1 kg dummy head holding neuronal cells and generating a ≤4,000 g-force acceleration upon impact. Dynamic impact led to both reduction in neuron viability and massive solubility changes in the proteome profiled using Proteome Integral Solubility Alteration (PISA) assay. The affected proteins mapped not only to the expected pathways like cell adhesion, collagen and laminin structures, as well as response to stress, but also to other dense protein networks, such as immune response, complement and coagulation cascades. The cellular effects are found to be mainly due to the shockwave rather than the g-force acceleration. Soft materials could reduce the impact severity only until being fully compressed. This study shows way to develop a proteome-based meter for measuring irreversible shockwave-induced cell damage and provides a resource for identifying TBI protein biomarkers and potential drug targets for developing products aiming at primary prevention and intervention.

## Introduction

Traumatic brain injury (TBI) is a serious global health problem ^1^, affecting around 70 million individuals every year ^2^. It is sometimes called the “silent epidemic”, because many cases are not registered and thus are not reflected in official statistics ^3,4^. Not only TBI victims but also their relatives suffer significantly, and the economic burden on the society is therefore very large ^5,6^. Over the last decades, several studies have found and confirmed that one of the more serious consequences of TBI is dementia, even in younger individuals ^7^. Recently, it was reported that 6.3% of TBI cases were associated with dementia within 15 years following an impact to the head ^8^. With an increasing interest in risky leisure activities worldwide, the numbers of dementia cases following TBI are expected to spike dramatically during the next decades. Although society in general has understood the advantages of primary prevention, such as using helmets, more efforts are required to reduce the severe consequences of TBI. Furthermore, conventional helmets do not always protect from severe consequences of head impact, as the well-known accident with Michael Schumacher has proven.

Today the treatment of moderate or severe TBI is still challenging. The most significant predictor of patients’ outcome is the development of cytotoxic brain tissue edema defined as a substantial increase in the intracellular water content leading to an expansion of brain volume ^9^. When severe enough, this causes an increased intracranial pressure usually refractory to existing osmotic treatments. The ultimate therapy choice is decompressive craniotomy which diminishes the intracranial hypertension ^10^. While this surgery significantly reduces the mortality rate, it creates further complications ^11,12^.

A number of molecular mechanisms underlying TBI are already known, but the molecular etiology of TBI and its effects at the cellular level are not completely understood. TBI involves the two stages of primary and secondary injury. While the primary injury occurs during direct insult to the brain, the secondary injury happens through subsequent biochemical changes such as ischemia, inflammation and cytotoxic processes ^13^. The main secondary TBI-induced injuries include Wallerian degeneration of axons defined as the disorganization of axonal cytoskeletal network ^14^, mitochondrial dysfunction ^13,15,16^, excitotoxicity by excessive release of excitatory amino acids ^17,18^, neuroinflammation through infiltration of circulating lymphocytes, neutrophils and monocytes into the injured brain parenchyma ^19^, oxidative stress ^15,20^ and consequent lipid peroxidation ^21^, impairment of autophagy and lysosomal pathways ^22–25^, as well as apoptotic cell death of neurons, oligodendrocytes and glia ^26–28^.

Extensive research has been conducted to identify druggable targets associated with these processes, to reduce and possibly prevent the adverse effects of TBI. However, most research was focused on the second-generation primary intracellular phenomena, such as excessive release of excitatory glutamate and aspartate from presynaptic nerve terminals ^17^, while first-generation primary processes occurring immediately after impact in brain cells have received significantly less attention. Potential therapeutics have been centered on stabilizing the site of injury and prevention of secondary damage. For example, glutamate receptor antagonists (for protection of neurons against excitotoxicity), antioxidants, antiinflammatory and anti-apoptotic agents, neurotrophic factors and stem cell therapies have been considered ^13^. However, our limited understanding of the TBI pathophysiology and the focus on secondary effects have hampered the development of effective TBI treatments. During the past three decades, more than 30 diagnostics or therapeutic pharmaceutical agents for TBI have failed in clinical trials ^13,29^. The primary injuries are believed to be hardly reversible ^13^; however, the history of biology and medicine knows examples of successful reversal of phenomena deemed irreversible by previous research; e.g., cell differentiation ^30^.

Therefore, in the current study, we focused on the first-generation primary molecular phenomena in TBI. Our working hypothesis was that external impact causes protein unfolding that triggers all subsequent phenomena in TBI. It is well known that high static pressure can unfold proteins ^31^. Previously we have shown by computer simulations and mechanistically simulated impact that dynamic pressure pulse can affect the aggregation state of laminin LN521, chosen as a representative protein due to its high abundance in the cells ^32^. To study such effects at the level of the whole proteome, here we employed a previously designed apparatus ^32^ and simulated dynamic impact on human brain-derived cells. The apparatus accommodates a cell culture plate with human neuroblastoma SH-SY5Y cells and allows for a 5.1 kg dummy head to fall from different heights on a static bottom plate (floor), inducing (depending on the height of impact) up to 4,000 g acceleration measured by computer-controlled sensor. To measure the effect of the impact on proteins’ aggregated state, we employed the recently introduced Proteome Integral Solubility Alteration (PISA) ^33^ assay, which is the high-throughput version of Thermal Proteome Profiling (TPP) ^34–36^. PISA can monitor the solubility changes occurring in the whole proteome upon a variety of cell-stimulating influences ^37–42^.

A dynamic impact of a dummy head holding the cell culture plate on a static bottom plate generates a pressure wave that is characterized by maximum acceleration (that can be measured in g’s) as well as the speed of propagation. The propagation speed is essential for the damage a wave creates. The initial wave caused by the impact, known as a shockwave ^43^, is supersonic in a given material and deposits energy in the material it passes through, causing damage at both molecular and supramolecular levels. A shockwave is responsible for, e.g., the well-known phenomenon of hard material erosion by water droplets falling on it with a speed as low as <3 m/s ^44^. After losing substantial energy, the shockwave slows down to the speed of sound, and the resulting acoustic wave propagates causing less damage at the molecular level, even though it may exert significant transient pressure. As we could measure only the acceleration g experienced by the cell culture but not the speed of the pressure wave passing through it, we had to untangle the effect on cells of the shockwave from the effect of acceleration.

## Results

### The dynamic impact experiments

The workflow of the dynamic impact experiment is shown in **Fig. 1**. The SH-SY5Y neuroblastoma cells were cultured in Nunclon dishes (diameter 35 mm; height 10 mm) to a density of 250,000 cells per dish, and the dish was placed into a 5.1 kg dummy head that was dropped from different heights (55-110 cm) on the static bottom plate (hard surface). In some experiments, the static bottom plate was covered with a mat composed of energy absorbing material (a rubber made of isobutylene and isoprene designed as pyramids of 6 mm length connected to a base membrane of 2 mm thickness). An ICP Shock accelerometer (PCB Piezotronics, SHEAR ICP shock accelerometer model 350D02 with sensitivity of ±1 g) was attached to the top of the dummy head. The PicoScope 6 software was used to record the g-force time diagram during the impact. Both impact-treated cells as well as control cells not exposed to the impact were then used for viability measurements and PISA analysis immediately after the impact ^33^. Alternatively, cells were kept in culture for 24-48 h for subsequent viability measurements. PISA analysis of the cells exposed to impact vs. control cells yielded information on protein’s changed solubility. As all experiments were performed in several replicates, statistical significance of each protein’s solubility change was calculated using two-tailed unpaired Student’s t-test. The most significantly changed proteins were mapped on known signaling and metabolic pathways.

**Fig. 1.**
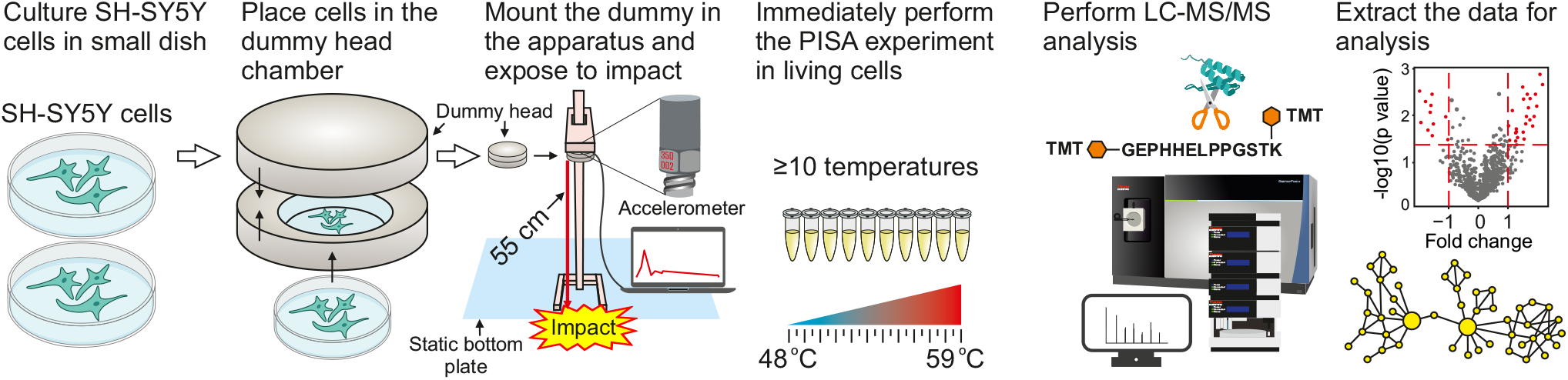
Workflow of dynamic impact simulation. See the text and methods for explanations. The dummy head holding neuronal cells falls on the static bottom plate (floor) with or without an energy absorbing material.

### Acceleration during impact

The g-force during impact was determined in a series composed of ≥4 independent measurements. On average, the impact from the 55 cm height on hard static bottom plate produced acceleration of 1,556 ± 456 g, while the impact from the 110 cm height produced acceleration of 3,959 ± 978 g. For the static bottom plate with the energy absorbing material, the corresponding values were 900 ± 289 g and 1,625 ± 510 g for 55 cm and 110 cm heights, respectively. A representative g-force time diagram for the 55 cm hard impact is shown in **Fig. 2a**.

**Fig. 2.**
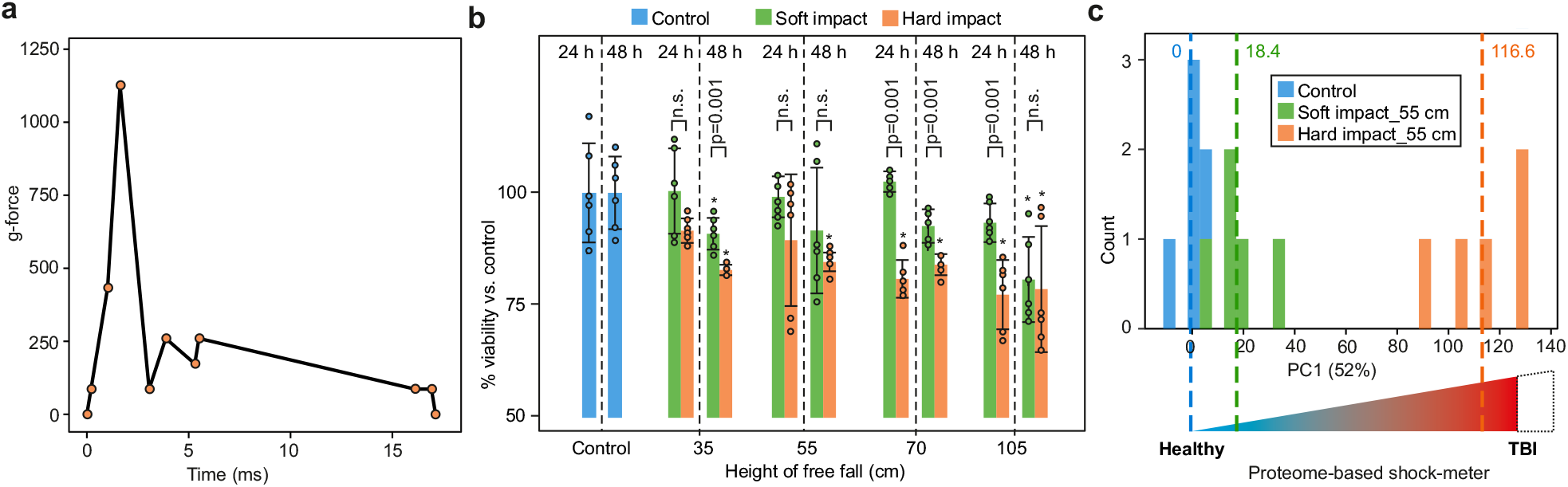
The effect of dynamic impact simulation on the viability and proteome of SH-SY5Y cell with and without energy absorbing mat. **a**, A representative g-force time diagram for 55 cm dynamic impact. **b**, Height dependent loss of cell viability in response to dynamic impact at 24 and 48 h. Protective effect of energy absorbing mat against loss of cell viability in response to dynamic impact at increasing heights (6 biological replicates; results shown as mean±SD; asterisks denote *p* value <0.05 vs. controls of the same timepoint; two-sided Student’s t-test). **c**, PCA component PC1 of PISA proteome signatures for controls and cells exposed to dynamic impact with and without energy absorbing mat (5 biological replicates). A conceptual schematic of the proteome-based shock-meter is shown.

### Dynamic impact reduces cell viability

After dynamic impact, the cells were kept in culture for 24 h and 48 h, after which measurements showed reduced cell viability in a height dependent manner compared to control at both time points. Notably, 48 h post hard impact the cell viability diminished compared to 24 h, indicating the persistence of cell death processes initiated by the impact (**Fig. 2b**). The energy absorbing mat significantly reduced the effect of dynamic impact on cell viability loss up to 70 cm height, but at 105 cm the viability after 48 h was statistically indistinguishable for the hard and soft impacts (**Fig. 2b**). The Trypan Blue dye exclusion assay showed no significant change in the percentage of dye-permeable cells upon impact, indicating that the cell membrane remained largely intact, and thus the cell viability loss was due to internal cell damage.

### Dynamic impact induces massive solubility changes in the cellular proteome

To investigate the mechanism of internal cell damage, the cells exposed to dynamic impact from 55 cm height on the hard surface, soft surface with energy absorbing material as well as the control cells were immediately processed after impact for PISA assay in 5 biological replicates. In total, 6,708 proteins were quantified by LC-MS/MS. After removing the contaminants and missing values, the final data contained 5,612 proteins (**Supplementary Data**).

At the first glance, 25% of the proteome (1,382 proteins) showed a significant change (at a *p* value cutoff of 0.05 and -0.5>log2 fold change>0.5) in solubility upon the hard impact, while only 1.5% of the proteome (86 proteins) showed a similar change upon impact on the plate covered with energy absorbing material. However, counting significant proteins is a less robust way of quantifying the proteome changes than using the coordinate in principal component analysis (PCA). Previously we have used such an approach for quantifying the transition of the cellular proteome from stemness to differentiation ^38^.

When the impact PISA data were subjected to PCA, the first principal component (PC1) accounted for the main part (52%) of data variation and separated the data points by the harshness of the impact, as expected (**Fig. 2c**). If the average position on PC1 of the control cells is taken as zero, the average coordinate of the cells after soft floor impact will be 18.4±13 and that of hard floor impact, 117±15. Therefore, PC1 of the PCA plot of PISA data can be used as a “shock-meter” for quantitatively assessing the effect of a given dynamic impact on the cells. However, later analysis showed the necessity to differentiate between the protein markers of the reversible and irreversible cell damage.

### Individual proteins changing solubility upon dynamic impact

On volcano plots in **Fig. 3a-b**, the proteins that changed their solubility significantly upon dynamic impact are shown in red. There were massive changes in protein solubility upon hard impact, which were alleviated to a large extent by the energy absorbing material.

**Fig. 3.**
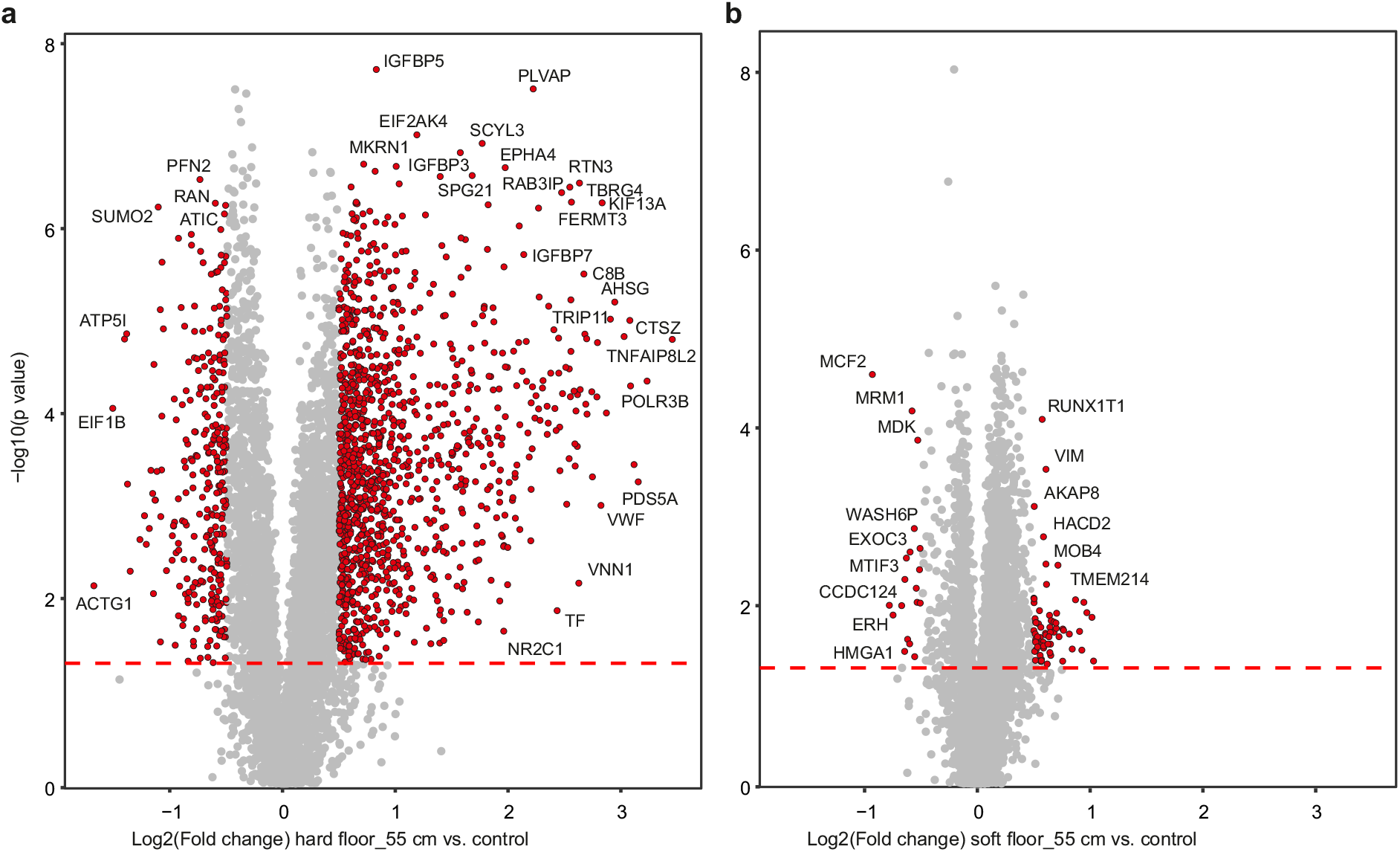
The extent of solubility changes across the proteome upon hard and soft impacts. Plotting the protein fold change (vs. control) against significance in PISA assay highlights the proteins with solubility/stability changes in response to dynamic impact on the hard (**a**) and soft surfaces (**b**) with a drop from 55 cm. The most important outliers (absolute log2 fold change vs. control >0.5 and *p* value <0.05) are highlighted (5 independent biological replicates).

The comparison between the hard and soft impacts on **Fig. 4a** shows that mostly the same proteins are affected in both cases, but the change in solubility is much larger upon hard than soft impact. Inspecting the proteins in **Fig. 4a** can be helpful in identification of potential diagnostic and/or therapeutic targets. Of particular interest is the group of proteins that changed their solubility in the same direction upon both hard and soft impacts, but with higher solubility change upon hard impact. Such proteins are highlighted in green and orange in **Fig. 4a** (the annotations can be found in **Supplementary Data**). The top protein in that group is tumor necrosis factor alpha-induced protein 8-like protein 2 or TNFAIP8L2 which is involved in maintaining immune homeostasis by preventing the hyperresponsiveness of the immune system ^45,46^. Among the top proteins of interest, collagens COL5A1, COL18A1 and COL21A1 have the laminin G domain (shown as orange dots), similar to laminin LN521 that has previously been found to unfold upon a dynamic impact, as assessed by electrophoresis and electron microscopy ^32^.

**Fig. 4.**
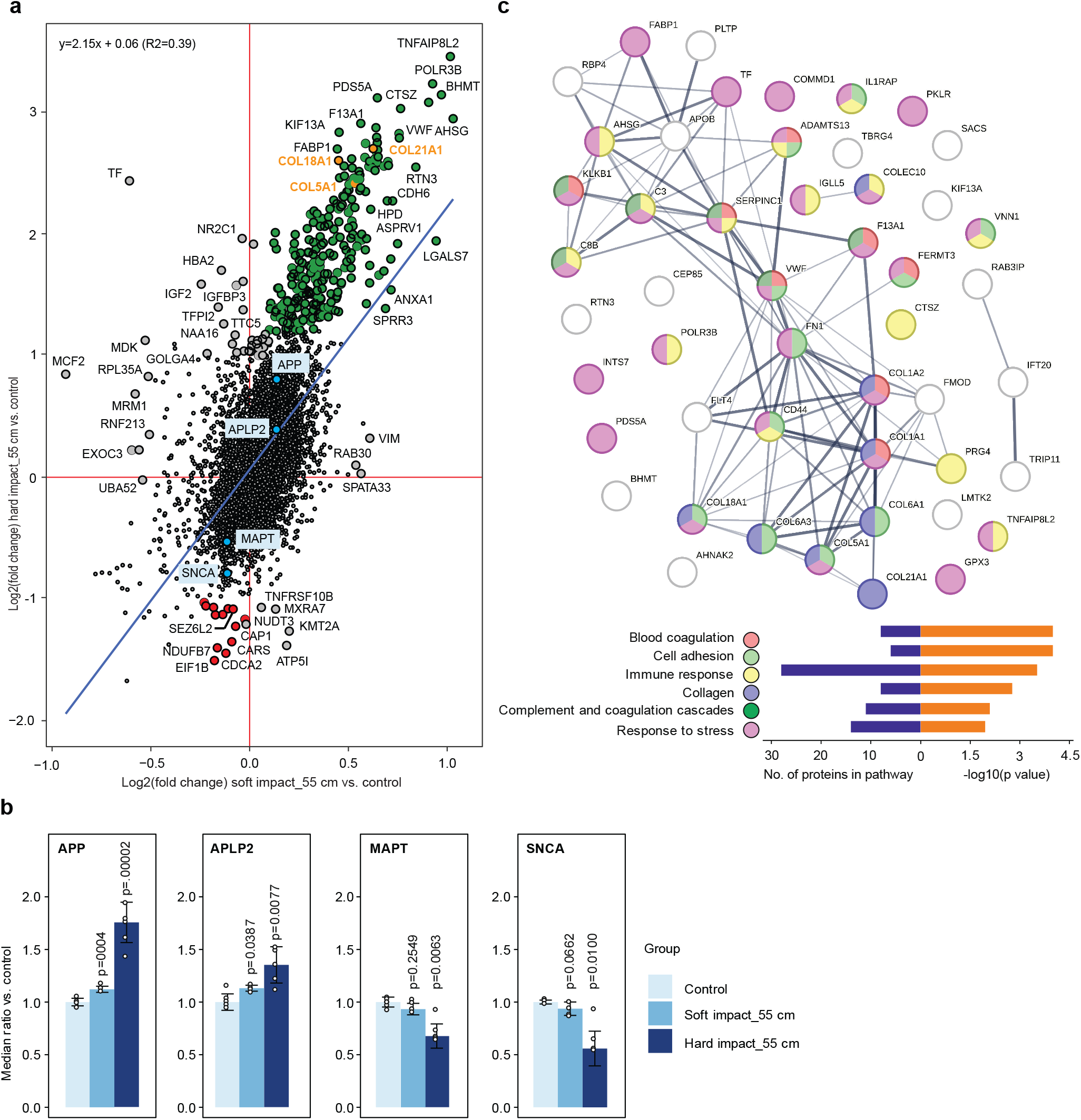
The massive proteome solubility changes affect essential cellular pathways. **a**, Comparing the proteome response under the two impact conditions. The proteins of special interest (see text) are shown in green (solubility increased), orange (collagens, solubility increased), red (solubility decreased) and sky blue (proteins known to be involved in the pathology of neurodegenerative diseases). **b**, Changes in the solubility of protein involved in the pathology of Alzheimer’s and Parkinson’s disease upon dynamic impact vs. control. **c**, Selected gene ontology pathways enriched for 50 top proteins (see text).

There was also a distinct group of proteins with reduced solubility (likely to become unfolded) upon dynamic impact (red dots in **Fig. 4a**; the annotations can be found in **Supplementary Data**). Four such proteins (i.e., ATP synthase subunit E or ATP5I, cytochrome b-c1 complex subunit 6 or UQCRH, NADH dehydrogenase [ubiquinone] 1 beta subcomplex subunit 7 or NDUFB7 and succinate dehydrogenase [ubiquinone] cytochrome b small subunit or SDHD) mapped to mitochondrial respirasome or electron transport change in the inner mitochondrial membrane (*p*=0.01). Interestingly, Seizure 6-like protein 2 or SEZ6L2 was also unfolded upon impact. SEZ6L2 contributes to specialized endoplasmic reticulum functions in neurons and is involved in regulation of neuritogenesis or neuron outgrowth ^47,48^. Autoantibodies against SEZ6L2 have been found in several patients with various forms of ataxia with atypical parkinsonism which usually occurs due to brain injury in regions controlling muscle coordination, such as the cerebellum ^49,50^.

### Dynamic impact modulates the solubility of proteins involved in neurodegenerative disorders

Neurodegenerative disorders such as Alzheimer’s and Parkinson’s diseases are associated with unfolding and aggregation of amyloid β and alpha-synuclein (SNCA) proteins, respectively. In our dynamic impact experiment, we observed solubility changes for amyloid precursor protein (APP), amyloid beta precursor like protein 2 (APLP2), microtubule-associated protein tau (MAPT) and SNCA (**Fig. 4b**). The solubility (or stability) of APP and APLP2 increased, while that of MAPT and SNCA decreased upon hard and soft dynamic impacts. Interestingly, the dynamic impact on the soft floor also changed the solubility of the above proteins in the same direction as the hard impact, however to a much lesser extent.

An interesting question is whether these solubility changes induce long-term consequences in neurons; if yes, this observation could link TBI to neurodegenerative disorders. Since neurons do not undergo cell division, they might accumulate such changes over lifetime. And given that the cellular repair systems such as autophagy are compromised in the above diseases ^51,52^, these solubility changes may contribute to or even trigger neuronal degeneration.

### Dynamic impact affects essential cellular processes

When top 50 proteins with highest significant changes upon hard impact were subjected to the network analysis in StringDB, they mapped not only to the expected pathways, such as cell adhesion, collagen trimer, and response to stress, but also to other dense protein networks, such as immune response, complement and coagulation cascades (**Fig. 4c**).

Coagulation factor XIII A chain (F13A1), von Willebrand factor (VWF), complement C3 beta chain (C3), antithrombin-III (SERPINC1), plasma kallikrein (KLKB1) and complement component C8 beta chain (C8B) that are involved in complement and coagulation cascades were among the proteins of interest in **Fig 4a**. The presence of immune response as well as complement and coagulation cascades among the affected pathways is noteworthy, and perhaps indicates that such proteins are evolutionarily designed to immediately respond to trauma, as there is not enough time for protein synthesis in response to imminent stress. The complement system is known to exert robust, immediate, and nonspecific immune responses ^53^. In single cell organisms, C3 provides constant surveillance of invading microbes by inducing protective inflammasome type responses ^54^. It has therefore been hypothesized that C3 is part of an evolutionary ancient system and is stored in vesicles, serving as an intracellular defense mechanism capable of quick release in defense against pathogenic invaders ^55^. C3 and other proteins involved in coagulation processes that are picked up in our analysis are apparently involved in sensing mechanical trauma. Such proteins may therefore be investigated as potential biomarkers of severe TBI. It should also be noted that some collagens such as COL1A1 and COL1A2 are involved in blood coagulation ^56^ and its activation ^57,58^, and can therefore trigger secondary damaging phenomena in TBI.

We also investigated the cellular components enriched among the proteins of interest. Among the most relevant components top 50 proteins mapped to were endoplasmic reticulum (17 proteins), extracellular matrix (16 proteins), secretory vesicle (13 proteins), Golgi apparatus (12 proteins), and cytoplasm (42 proteins) (all *p* values < 0.02). These results may contain clues on the mechanism of primary cellular damage upon dynamic impact.

### The cellular effects are mainly due to the impact shockwaves

To investigate whether the cellular effects upon dynamic impact are mostly due to g-force acceleration or shockwaves, SH-SY5Y cells suspended in PBS were exposed for 10 min to centrifugal force equivalent to that experienced upon dynamic impact, after which cells were plated back and kept in culture, with cell viability measured after 24 h and 48 h. At centrifugal g-forces (around 1,250) similar to those experienced in hard dynamic impact at 55 cm and soft impact at 110 cm, the reduction in cell viability after 24 h was also similar (**Fig. 5a**). However, the cell viability did not decrease significantly even at 20,000 g centrifugal acceleration and, remarkably, the viability at 48 h increased (vs. controls) compared to 24 h at all centrifugal forces, indicating that at least part of the cellular damage was reversible. Note also that the dynamic impact occurs on the millisecond time scale while the centrifugation was performed over 10 min.

**Fig. 5.**
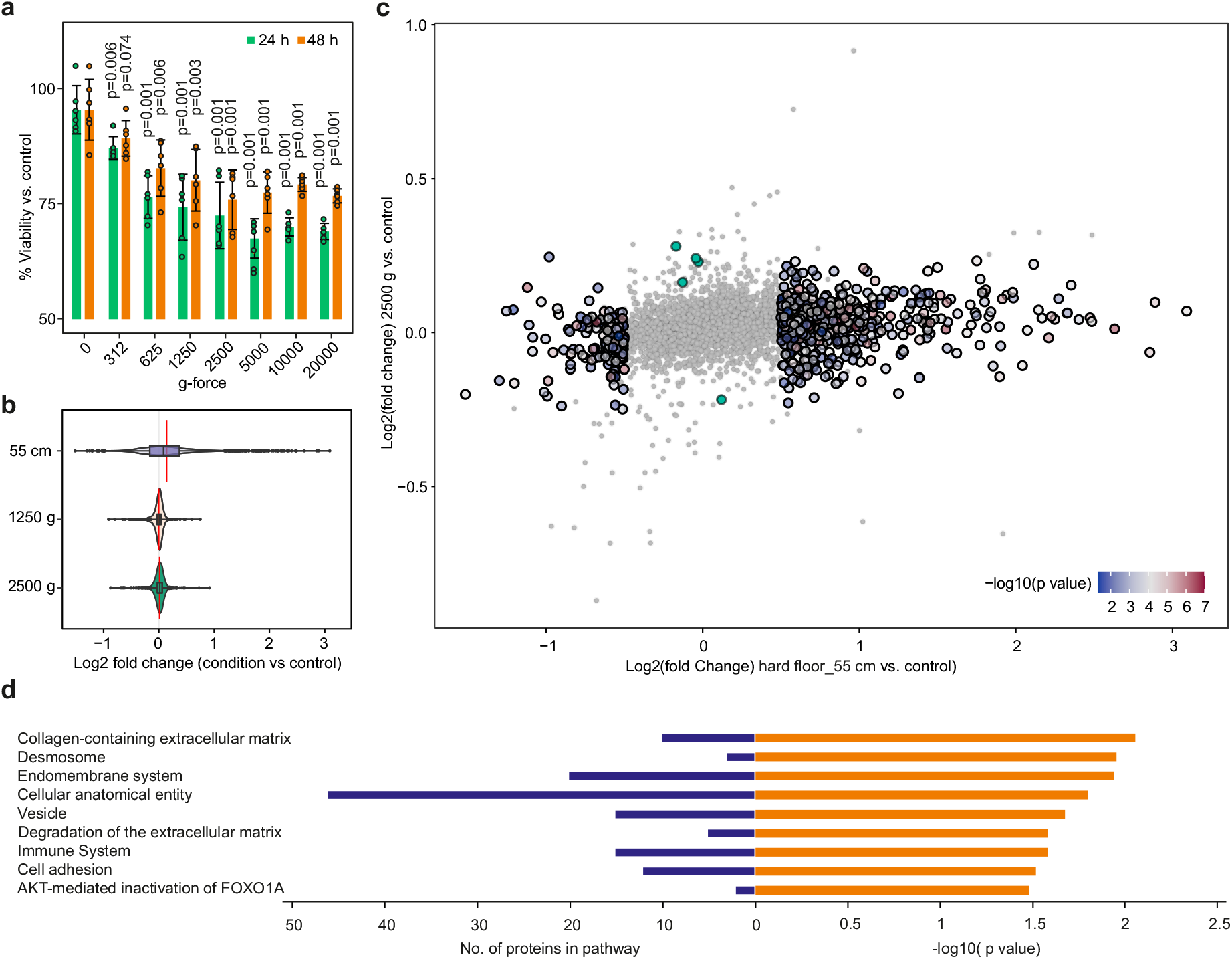
Exposing cells to centrifugal g-force acceleration reduces cell viability in a partially reversible manner and induces only minimal changes in the solubility of the proteome. **a**, Reduction of cell viability upon centrifugation of SH-SY5Y cells in PBS for 10 min at different g-forces (6 biological replicates; *p* values between the given g-force vs. control; two-sided Student’s t-test). **b**, The extent of overall proteome solubility change in response to the hard impact (55 cm) vs. centrifugation for 10 min at 1250 and 2500 g. **c**, Solubility changes in response to the hard impact (55 cm) vs. centrifugation for 10 min at 2500 g. **d**, The enriched pathways and processes for the top 50 proteins only changing significantly in response to dynamic hard impact (and not centrifugation at 2500 g).

To separate the effect of acceleration vs. shockwaves on the cell proteome, we subjected the centrifuged cells to PISA analysis, merging the datasets for 1,250 and 2,500 g and comparing them with PISA data from dynamic impact at 55 cm on the hard floor. The changes in proteome solubility upon dynamic impact were drastically larger compared to those in cells exposed only to comparable accelerations (**Fig. 5b-c**). **Fig. 5d** shows pathways that are specifically enriched for the set of top 50 proteins changing in response to dynamic impact and not centrifugation. These included expected pathways such as cell adhesion and immune system, as well as cellular anatomical features.

On the other hand, only 5 proteins showed a significant change in solubility upon centrifugation at 2500 g, with no change upon dynamic impact from 55 cm on the hard floor. These proteins did not map to any specific pathways.

## Discussion

Here we demonstrate a new methodology for simulation of TBI using cultured neuronal cells. Exposing cells to centrifugal g-forces had a similar effect on cell survival after 24 h as a dynamic impact with an equivalent g-force, even though a much larger magnitude of the effect was expected given the 6 orders of magnitude difference in treatment time scales (10 min vs. few milliseconds). Also, in centrifugal acceleration experiments, the cells partially recovered after 48 h, while after dynamic impact their viability decreased further over time. The energy absorbing material could partially rescue the cellular metabolic viability, but not for the most severe impacts. Experimental as well theoretical studies showed that soft materials can reduce the impact severity only until being fully compressed, and upon reaching that point they can transmit shockwaves nearly as easy as hard materials ^44^.

Taken together, these observations indicate that most of the irreversible phenomena in cells are the result of shockwaves emanating from the dynamic impact, while the g-force acceleration plays a less prominent role in irreversible cellular damage. As the Trypan Blue dye exclusion assay results revealed that the cell membrane remained largely intact even upon the hardest impact, the cell viability loss must be due to internal cell damage.

By PISA profiling we found that the damage induced by dynamic impact is likely to relate to the solubility changes of many cellular proteins, including those involved in the pathology of neurodegenerative diseases. This supported our previous findings on alteration of laminin structure by dynamic impact ^32^. As most of the detected solubility changes were positive (green and orange dots in **Fig. 4a**), disruption of protein complexes or organelles seems to be the most likely damaging effect, while protein unfolding (red dots in **Fig. 4a**) exacerbated the damage. This conclusion was supported by pathway analysis of most affected proteins that changed their solubility proportionally to impact hardness.

The overall effect on proteome solubility is well represented by PC1 in PCA. These results open the way for developing a proteome-based shock-meter, as conceptually shown in **Fig. 2c**. Such a shock-meter can be used to monitor the relative effect of dynamic impacts on neural cells under various circumstances, and help designing more effective preventive measures such as helmets and flooring materials. Cerebral concussion after kinetic energy impact is the most common complication often defined as mild TBI. At the same time this is the most puzzling condition in many cases characterized by different clinical symptoms ^61^. Concussion currently has no generally accepted theoretical definition, as it cannot be monitored using the available imaging technologies. Thus, different kinds of TBI can be theoretically graded using a proteome-based shock-meter.

Low mechanical dynamic and static pressure forces can both influence structure of proteins and protein complexes ^62^. The protein tertiary and quaternary structures rely on hydrophobic interactions and hydrogen bonds to maintain their three-dimensional configuration for proper protein functioning. When these bonds are disrupted, e.g. under pressure or mechanical shock, the proteins can undergo unfolding events that may be followed by misfolding ^32,63^. Unfolded proteins usually lose their solubility, which can be measured by PISA assay. While a distinct group of protein seemed to unfold upon dynamic impact, most proteins increased their solubility (**Figs. 3a** and **4a**), which likely resulted from breakage of the complexes they are involved in. A scenario is also possible where a protein in a complex unfolds upon the impact, which disrupts the complex and releases from it several proteins, thus increasing their solubility.

Soluble proteins are enshrined in a hydration shell that is often more dense than bulk water ^64^. It is thus feasible that, as many proteins become more soluble upon the dynamic impact, the overall volume of the hydration shells expands, necessitating water uptake from the extracellular and interstitial area, resulting in swelling of the cells in TBI and eventuating in cytotoxic brain tissue edema.

Upon the breakage of the protein complex, the proteins may refold and form the complex again – in this case, the effect is reversible. We observed such an effect in centrifugal acceleration experiments. Refolded proteins would not show marked solubility change in a PISA analysis. On the other hand, irreversible damage to proteins observed by PISA assay upon hard impact is likely to be responsible for triggering the cell death pathways responsible for continuous reduction in cellular viability between 24 h and 48 h post impact. The PISA-based proteome-based shock-meter is likely to be reflective of the irreversible cellular damage. Thus, using such a shock-meter can help test and design energy absorbing materials that reduce or prevent irreversible cell damage. Furthermore, the data presented in the current study will serve as a resource for follow-up studies on the acute and prolonged molecular consequences of massive proteome unfolding in TBI and potentially its link to increased incidence of dementia and potentially other neurodegenerative disorders. Finally, the described proteome approaches may help in development of innovative products for primary prevention of TBI by providing protein targets for intervention as well as a means for assessing the preventive effect at the molecular level.

## Materials and methods

### Cell culture

Human SH-SY5Y cells (ATCC, USA) were grown at 37°C in 5% CO2 using DMEM/F12 medium with GlutaMAX (cat#31331093, Thermo) supplemented with 10% heat-inactivated FBS (cat#A3840402, Thermo), 100 units/mL penicillin/streptomycin (cat#15070063, Thermo), MEM non-essential amino acids (cat#11140035) and HEPES buffer (cat#15630056, Thermo). Low-number passages were used for the experiments.

### Simulation of dynamic impact at different heights

Cells were seeded at a density of 200,000 per well in Nunclon cell culture dishes (diameter 35 mm × 10 mm) and grown for 24 h. The dishes were then loaded into dummy head of 5.1 kg weight and the impact experiment was performed at different heights in 5 biological replicates. Cells not exposed to impact were used as controls (controls were kept on the bench for the same duration of the dynamic impact experiment to minimize the effect of other variables). Depending on the experiment, cells underwent processing immediately or were kept in culture for different time points before processing (the media volume was restored after the impact, but the detached cells were kept in culture).

### Cell viability measurements

Cells were seeded at a density of 200,000 per well in Nunclon cell culture dishes and grown for 24 h. The impact experiment was performed similar to above by dropping the dummy head from different heights. The control and treated samples were kept for 24 and 48 h. The cells detached upon the impact were kept in the dishes. Thereafter cell viability was measured using CellTiter-Blue Assay (Promega) according to the manufacturer protocol, as described previously ^65^.

For measuring the viability of centrifuged cells, 250,000 cells were aliquoted in 300 μL PBS and centrifuged at 312, 625, 1250, 2500, 5000, 10,000 and 20,000 g for 10 min. Thereafter, PBS was removed, fresh media were added, and cells were cultured in 6-well plates for 24 and 48 h. The viability measurement was performed like above.

### PISA assay

Cells were processed according to PISA assay protocol ^33^. After the impact, the media were removed, and cells were washed with PBS. 350 μL of PBS was added to each plate, after which cells were scraped from the surface. Each sample was then divided into 10 aliquots in PCR plates. In the case of centrifugation experiment, samples in PBS were directly aliquoted into the PCR plates after centrifugation. The plate was heated in an Eppendorf gradient thermocycler (Mastercycler X50s) at the temperature range of 48-59°C for 3 min. Samples were cooled for 3 min at room temperature and afterwards snap frozen in liquid nitrogen. Samples from each replicate were then combined. The samples were freeze-thawed in liquid nitrogen twice more, and then transferred into polycarbonate thick-wall tubes and centrifuged at 100,000 g and 4°C for 20 min.

The soluble protein fraction was transferred to new Eppendorf tubes. Protein concentration was measured in all samples using Pierce BCA Protein Assay Kit (Thermo), the volume corresponding to 25 μg of protein was transferred from each sample to new tubes and urea was added to a final concentration of 4 M. Dithiothreitol (DTT) was added to a final concentration of 10 mM and samples were incubated for 1 h at room temperature. Subsequently, iodoacetamide (IAA) was added to a final concentration of 50 mM and samples were incubated at room temperature for 1 h in the dark. The reaction was quenched by adding an additional 10 mM of DTT. The proteins were then precipitated using methanol/chloroform, and after drying, resuspended in 20 mM EPPS pH 8.5 with 8 M urea. Then, urea was diluted to 4 M with 20 mM EPPS and then Lysyl endopeptidase (LysC; Wako) was added at a 1:50 w/w ratio at room temperature overnight. Samples were diluted with 20 mM EPPS to the final urea concentration of 1 M, and trypsin was added at a 1:50 w/w ratio, followed by incubation for 6 h at room temperature. Acetonitrile (ACN) was added to a final concentration of 20% and TMTpro-16plex reagents were added 4x by weight (200 μg) to each sample, followed by incubation for 2 h at room temperature. The reaction was quenched by addition of 0.5% hydroxylamine. Samples within each replicate were combined, acidified by Trifluoroacetic acid (TFA), cleaned using Sep-Pak cartridges (Waters) and dried using DNA 120 concentrator (Thermo). The pooled samples were resuspended in 20 mM ammonium hydroxide and separated into 96 fractions on an XBrigde BEH C18 2.1×150 mm column (Waters; Cat#186003023), using a Dionex Ultimate 3000 2DLC system (Thermo Scientific) over a 48 min gradient of 1-63% B (B=20 mM ammonium hydroxide in acetonitrile) in three steps (1-23.5% B in 42 min, 23.5-54% B in 4 min and then 54-63% B in 2 min) at 200 μL min^-1^ flow. Fractions were then concatenated into 16 samples in sequential order. PISA samples subjected to acceleration in the centrifuge were concatenated into 12 samples in sequential order after fractionation.

### LC-MS/MS

After drying, samples were dissolved in buffer A (0.1% formic acid and 2% ACN in water). The samples were loaded onto a 50 cm EASY-Spray column (75 μm internal diameter, packed with PepMap C18, 2 μm beads, 100 Å pore size) connected to a nanoflow Dionex UltiMate 3000 UHPLC system (Thermo) and eluted in an organic solvent gradient increasing from 4% to 26% (B: 98% ACN, 0.1% FA, 2% H2O) at a flow rate of 300 nL min^-1^ over 180 min. The eluent was ionized by electrospray and mass spectra of the molecular ions were acquired with an Orbitrap Q Exactive HF mass spectrometer (Thermo Fisher Scientific) in data-dependent mode at MS1 resolution of 120,000 and MS2 resolution of 60,000, in the m/z range from 375 to 1500. Peptide fragmentation was performed via higher-energy collision dissociation (HCD) with energy set at 35% NCE. PISA samples subjected to acceleration in the centrifuge were analyzed using similar settings over a total 150 min gradient.

### Data processing

The raw LC-MS data were analyzed by MaxQuant version 1.6.2.3 ^66^. The Andromeda search engine ^67^ matched MS/MS data against the UniProt complete proteome database (human, version UP000005640_9606, 92,957 entries), unless otherwise specified. Trypsin/P was selected as enzyme specificity. No more than two missed cleavages were allowed. A 1% false discovery rate was used as a filter at both protein and peptide levels. For all other parameters, the default settings were used. After removing all the contaminants, only proteins with at least two peptides were included in the final dataset. Protein intensities were normalized by total abundance in each TMT channel and then Log2 transformed.

### Network mapping

For pathway analyses, STRING version 11.5 protein network analysis tool was used with default parameters ^68^.

### Statistical Analysis

Data analysis was performed using R project versions 3.6.1. Analysis of significance was performed with two-sided Student’s t-test throughout the manuscript.

## Data availability

The authors declare that all data supporting the findings of this study are available within the paper and its supplementary information files. All relevant data are available from the corresponding authors (H.v.H. and R.A.Z.). The mass spectrometry data that support the findings of this study have been deposited in ProteomeXchange Consortium (https://www.ebi.ac.uk/pride/) via the PRIDE partner repository ^69^ with the dataset identifier PXD037675 (impact experiment) and PXD038607 (g experiment).

## Acknowledgements

This research is funded by grants from KI SFO awarded to R.A.Z.; A.A.S. was supported by Swedish Research Council (grant 2020-00687) and the Swedish Society of Medicine (grant SLS-961262, 1086 Stiftelsen Albert Nilssons forskningsfond).

## Author contributions

Conceptualization, H.v.H., A.A.S., and R.A.Z.; methodology and experiment design, A.A.S., H.v.H., H.G., and R.A.Z.; project organization, resources and funding acquisition, H.v.H. and R.A.Z.; PISA experiments, A.A.S., H.L., B.N., M.J., and H.G.; data analysis and visualization, H.G. and A.A.S; writing—review & editing, A.A.S., R.A.Z. and H.v.H.

## Conflicts of interest

Authors declare no conflicts of interest.

## Notes

### Competing Interest Statement

The authors have declared no competing interest.

## References

1. von Holst, H. Traumatic Brain Injury in Handbook of clinical neuroepidemiology. (Nova Publishers, 2007).

2. Dewan, M. C. et al. Estimating the global incidence of traumatic brain injury. J. Neurosurg. (2019) doi:10.3171/2017.10.JNS17352.

3. Rusnak, M. Traumatic brain injury: Giving voice to a silent epidemic. Nat. Rev. Neurol. (2013) doi:10.1038/nrneurol.2013.38.

4. Goldstein, M. Traumatic brain injury: A silent epidemic. Annals of Neurology (1990) doi:10.1002/ana.410270315.

5. Perlesz, A., Kinsella, G. & Crowe, S. Impact of traumatic brain injury on the family: A critical review. Rehabilitation Psychology (1999) doi:10.1037/0090-5550.44.1.6.

6. Degeneffe, C. E. Family caregiving and traumatic brain injury. Heal. Soc. Work (2001) doi:10.1093/hsw/26.4.257.

7. Gardner, R. C. et al. Dementia risk after traumatic brain injury vs nonbrain trauma: The role of age and severity. JAMA Neurol. (2014) doi:10.1001/jamaneurol.2014.2668.

8. Nordström, A. & Nordström, P. Traumatic brain injury and the risk of dementia diagnosis: A nationwide cohort study. PLoS Med. (2018) doi:10.1371/journal.pmed.1002496.

9. Unterberg, A. W., Stover, J., Kress, B. & Kiening, K. L. Edema and brain trauma. Neuroscience (2004) doi:10.1016/j.neuroscience.2004.06.046.

10. Kolias, A. G., Kirkpatrick, P. J. & Hutchinson, P. J. Decompressive craniectomy: Past, present and future. Nature Reviews Neurology (2013) doi:10.1038/nrneurol.2013.106.

11. Stiver, S. I. Complications of decompressive craniectomy for traumatic brain injury. Neurosurgical Focus (2009) doi:10.3171/2009.4.FOCUS0965.

12. Honeybul, S. & Ho, K. M. Long-term complications of decompressive craniectomy for head injury. J. Neurotrauma (2011) doi:10.1089/neu.2010.1612.

13. Ng, S. Y. & Lee, A. Y. W. Traumatic Brain Injuries: Pathophysiology and Potential Therapeutic Targets. Frontiers in Cellular Neuroscience (2019) doi:10.3389/fncel.2019.00528.

14. Tang-Schomer, M. D., Patel, A. R., Baas, P. W. & Smith, D. H. Mechanical breaking of microtubules in axons during dynamic stretch injury underlies delayed elasticity, microtubule disassembly, and axon degeneration. FASEB J. (2010) doi:10.1096/fj.09-142844.

15. Xiong, Y., Gu, Q., Peterson, P. L., Muizelaar, J. P. & Lee, C. P. Mitochondrial dysfunction and calcium perturbation induced by traumatic brain injury. J. Neurotrauma (1997) doi:10.1089/neu.1997.14.23.

16. Singh, I. N., Sullivan, P. G., Deng, Y., Mbye, L. H. & Hall, E. D. Time course of post-traumatic mitochondrial oxidative damage and dysfunction in a mouse model of focal traumatic brain injury: Implications for neuroprotective therapy. J. Cereb. Blood Flow Metab. (2006) doi:10.1038/sj.jcbfm.9600297.

17. Chamoun, R., Suki, D., Gopinath, S. P., Goodman, J. C. & Robertson, C. Role of extracellular glutamate measured by cerebral microdialysis in severe traumatic brain injury: Clinical article. J. Neurosurg. (2010) doi:10.3171/2009.12.JNS09689.

18. Faden, A. I., Demediuk, P., Panter, S. S. & Vink, R. The role of excitatory amino acids and NMDA receptors in traumatic brain injury. Science (80-.). (1989) doi:10.1126/science.2567056.

19. Lotocki, G. et al. Alterations in blood-brain barrier permeability to large and small molecules and leukocyte accumulation after traumatic brain injury: Effects of post-traumatic hypothermia. J. Neurotrauma (2009) doi:10.1089/neu.2008.0802.

20. Shohami, E. & Kohen, R. The Role of Reactive Oxygen Species in the Pathogenesis of Traumatic Brain Injury. in Oxidative Stress and Free Radical Damage in Neurology (2011). doi:10.1007/978-1-60327-514-9_7.

21. Ansari, M. A., Roberts, K. N. & Scheff, S. W. Oxidative stress and modification of synaptic proteins in hippocampus after traumatic brain injury. Free Radic. Biol. Med. (2008) doi:10.1016/j.freeradbiomed.2008.04.038.

22. Diskin, T. et al. Closed head injury induces upregulation of beclin 1 at the cortical site of injury. J. Neurotrauma (2005) doi:10.1089/neu.2005.22.750.

23. Clark, R. S. B. et al. Autophagy is increased in mice after traumatic brain injury and is detectable in human brain after trauma and critical illness. Autophagy (2008) doi:10.4161/auto.5173.

24. Sakai, K., Fukuda, T. & Iwadate, K. Immunohistochemical analysis of the ubiquitin proteasome system and autophagy lysosome system induced after traumatic intracranial injury: Association with time between the injury and death. Am. J. Forensic Med. Pathol. (2014) doi:10.1097/PAF.0000000000000067.

25. Au, A. K. et al. Autophagy Biomarkers Beclin 1 and p62 are Increased in Cerebrospinal Fluid after Traumatic Brain Injury. Neurocrit. Care (2017) doi:10.1007/s12028-016-0351-x.

26. Beer, R. et al. Temporal profile and cell subtype distribution of activated caspase-3 following experimental traumatic brain injury. J. Neurochem. (2000) doi:10.1046/j.1471-4159.2000.0751264.x.

27. Grady, M. S., Charleston, J. S., Maris, D., Witgen, B. M. & Lifshitz, J. Neuronal and Glial Cell Number in the Hippocampus after Experimental Traumatic Brain Injury: Analysis by Stereological Estimation. J. Neurotrauma (2003) doi:10.1089/089771503770195786.

28. Smith, D. H. et al. Progressive atrophy and neuron death for one year following brain trauma in the rat. J. Neurotrauma (1997) doi:10.1089/neu.1997.14.715.

29. Veenith, T. & Goon, S. S. Molecular mechanisms of traumatic brain injury: The missing link in management. World J. Emerg. Surg. (2009) doi:10.1186/1749-7922-4-4.

30. Takahashi, K. & Yamanaka, S. Induction of Pluripotent Stem Cells from Mouse Embryonic and Adult Fibroblast Cultures by Defined Factors. Cell (2006) doi:10.1016/j.cell.2006.07.024.

31. Kim, Y. S., Randolph, T. W., Seefeldt, M. B. & Carpenter, J. F. High-Pressure Studies on Protein Aggregates and Amyloid Fibrils. Methods in Enzymology (2006) doi:10.1016/S0076-6879(06)13013-X.

32. von Holst, H., Purhonen, P., Lanner, D., Balakrishnan Kumar, R. & Hebert, H. White Shark Protein Metabolism may be a Model to Improve the Outcome of Cytotoxic Brain Tissue Edema and Cognitive Deficiency after Traumatic Brain Injury and Stroke. J. Neurol. Neurobiol. 4, (2018).

33. Gaetani, M. et al. Proteome Integral Solubility Alteration: A High-Throughput Proteomics Assay for Target Deconvolution. J. Proteome Res. (2019) doi:10.1021/acs.jproteome.9b00500.

34. Savitski, M. M. et al. Tracking cancer drugs in living cells by thermal profiling of the proteome. Science (80-.). (2014) doi:10.1126/science.1255784.

35. Molina, D. M. et al. Monitoring drug target engagement in cells and tissues using the cellular thermal shift assay. Science (80-.). (2013) doi:10.1126/science.1233606.

36. Saei, A. A. et al. Comprehensive chemical proteomics for target deconvolution of the redox active drug auranofin. Redox Biol. 101491 (2020).

37. Saei, A. A. et al. System-wide identification and prioritization of enzyme substrates by thermal analysis. Nat. Commun. (2021).

38. Sabatier, P. et al. An integrative proteomics method identifies a regulator of translation during stem cell maintenance and differentiation. Nat. Commun. (2021) doi:10.1038/s41467-021-26879-4.

39. Becher, I. et al. Pervasive Protein Thermal Stability Variation during the Cell Cycle. Cell (2018) doi:10.1016/j.cell.2018.03.053.

40. Dai, L. et al. Modulation of Protein-Interaction States through the Cell Cycle. Cell (2018) doi:10.1016/j.cell.2018.03.065.

41. Potel, C. M. et al. Impact of phosphorylation on thermal stability of proteins. bioRxiv (2020). doi:10.1101/2020.01.14.903849.

42. Saei, A. A. et al. Mapping the GALNT1 substrate landscape with versatile proteomics tools. bioRxiv (2022).

43. Courant, R. & Friedrichs, K. O. Supersonic flow and shock waves. vol. 21 (Springer Science & Business Media, 1999).

44. Sun, T.-P. et al. Stress distribution and surface shock wave of drop impact. Nat. Commun. 13, 1–8 (2022).

45. Sun, H. et al. TIPE2, a Negative Regulator of Innate and Adaptive Immunity that Maintains Immune Homeostasis. Cell (2008) doi:10.1016/j.cell.2008.03.026.

46. Zhang, Y. et al. TIPE2, a novel regulator of immunity, protects against experimental stroke. J. Biol. Chem. (2012) doi:10.1074/jbc.M112.348755.

47. Yaguchi, H. et al. Sez6l2 regulates phosphorylation of ADD and neuritogenesis. Biochem. Biophys. Res. Commun. (2017) doi:10.1016/j.bbrc.2017.10.047.

48. Boonen, M. et al. Cathepsin D and its newly identified transport receptor SEZ6L2 can modulate neurite outgrowth. J. Cell Sci. (2016) doi:10.1242/jcs.179374.

49. Yaguchi, H. et al. Identification of anti-Sez6l2 antibody in a patient with cerebellar ataxia and retinopathy. Journal of Neurology (2014) doi:10.1007/s00415-013-7134-5.

50. Borsche, M. et al. Sez6l2-antibody-associated progressive cerebellar ataxia: a differential diagnosis of atypical parkinsonism. Journal of Neurology (2019) doi:10.1007/s00415-018-9115-1.

51. Nixon, R. A. & Yang, D. S. Autophagy failure in Alzheimer’s disease-locating the primary defect. Neurobiology of Disease (2011) doi:10.1016/j.nbd.2011.01.021.

52. Hou, X., Watzlawik, J. O., Fiesel, F. C. & Springer, W. Autophagy in Parkinson’s Disease. Journal of Molecular Biology (2020) doi:10.1016/j.jmb.2020.01.037.

53. Dunkelberger, J. R. & Song, W. C. Complement and its role in innate and adaptive immune responses. Cell Res. (2010) doi:10.1038/cr.2009.139.

54. Tam, J. C. H., Bidgood, S. R., McEwan, W. A. & James, L. C. Intracellular sensing of complement C3 activates cell autonomous immunity. Science (80-.). (2014) doi:10.1126/science.1256070.

55. Elvington, M., Liszewski, M. K. & Atkinson, J. P. Evolution of the complement system: from defense of the single cell to guardian of the intravascular space. Immunological Reviews (2016) doi:10.1111/imr.12474.

56. Mosher, D. F., Schad, P. E. & Kleinman, H. K. Cross-linking of fibronectin to collagen by blood coagulation factor XIIIa. J. Clin. Invest. (1979) doi:10.1172/JCI109524.

57. Zillmann, A. et al. Platelet-associated tissue factor contributes to the collagen-triggered activation of blood coagulation. Biochem. Biophys. Res. Commun. (2001) doi:10.1006/bbrc.2001.4399.

58. Niewiarowski, S., Stuart, R. K., Thomas, D. P. & Chalmers, T. C. Activation of Intravascular Coagulation by Collagen. Proc. Soc. Exp. Biol. Med. (1966) doi:10.3181/00379727-123-31440.

59. Batra, N. et al. Direct regulation of osteocytic connexin 43 hemichannels through AKT kinase activated by mechanical stimulation. J. Biol. Chem. (2014) doi:10.1074/jbc.M114.550608.

60. Liang, Y. et al. Meta-Analysis-Assisted Detection of Gravity-Sensitive Genes in Human Vascular Endothelial Cells. Front. Cell Dev. Biol. (2021) doi:10.3389/fcell.2021.689662.

61. Mayer, A. R., Quinn, D. K. & Master, C. L. The spectrum of mild traumatic brain injury: a review. Neurology 89, 623–632 (2017).

62. Gregoire, S., Irwin, J. & Kwon, I. Techniques for monitoring protein misfolding and aggregation in vitro and in living cells. Korean Journal of Chemical Engineering (2012) doi:10.1007/s11814-012-0060-x.

63. Roche, J. & Royer, C. A. Lessons from pressure denaturation of proteins. Journal of the Royal Society Interface (2018) doi:10.1098/rsif.2018.0244.

64. Kuffel, A. & Zielkiewicz, J. Why the solvation water around proteins is more dense than bulk water. J. Phys. Chem. B (2012) doi:10.1021/jp305172t.

65. Saei, A. A. et al. ProTargetMiner as a proteome signature library of anticancer molecules for functional discovery. Nat. Commun. (2019) doi:10.1038/s41467-019-13582-8.

66. Cox, J. & Mann, M. MaxQuant enables high peptide identification rates, individualized p.p.b.-range mass accuracies and proteome-wide protein quantification. Nat. Biotechnol. (2008) doi:10.1038/nbt.1511.

67. Cox, J. et al. Andromeda: A peptide search engine integrated into the MaxQuant environment. J. Proteome Res. (2011) doi:10.1021/pr101065j.

68. Szklarczyk, D. et al. The STRING database in 2017: quality-controlled protein–protein association networks, made broadly accessible. Nucleic Acids Res. gkw937 (2016).

69. Vizcaíno, J. A. et al. ProteomeXchange provides globally coordinated proteomics data submission and dissemination. Nature Biotechnology (2014) doi:10.1038/nbt.2839.

